# Zebrafish (*Danio rerio*) behavioral phenotypes not underscored by different gut microbiota

**DOI:** 10.1101/2024.05.29.596447

**Authors:** Paul A. Ayayee, Ryan Y. Wong

**Affiliations:** Department of Biology, University of Nebraska at Omaha, Omaha, NE, USA

**Keywords:** Zebrafish, Behavioral phenotype, microbiome, bold, shy

## Abstract

Different animal behavioral phenotypes maintained and selectively bred over multiple generations may be underscored by dissimilar gut microbial community compositions or not have any significant dissimilarity in community composition. Operating within the microbiota-gut-brain axis framework, we anticipated differences in gut microbiome profiles between zebrafish (*Danio rerio*) selectively bred to display the bold and shy personality types. This would highlight gut microbe-mediated effects on host behavior. To this end, we amplified and sequenced a fragment of the 16S rRNA gene from the guts of bold and shy zebrafish individuals (n=10) via Miseq. We uncovered no significant difference in within-group microbial diversity nor between-group microbial community composition of the two behavioral phenotypes. Interestingly, though not statistically different, we determined that the gut microbial community of the bold phenotype was dominated by *Burkholderiaceae, Micropepsaceae,* and *Propionibacteriaceae*. In contrast, the shy phenotype was dominated by *Beijerinckaceae, Pirelullacaeae, Rhizobiales_Incertis_Sedis*, and *Rubinishaeraceae.* The absence of any significant difference in gut microbiota profiles between the two phenotypes would suggest that in this species, there might exist a stable “core” gut microbiome, regardless of behavioral phenotypes, and or possibly, a limited role for the gut microbiota in modulating this selected-for host behavior. This is the first study to characterize the gut microbial community of distinct innate behavioral phenotypes of the zebrafish (that are not considered dysbiotic states) and not rely on antibiotic or probiotic treatments to induce changes in behavior. Such studies are crucial to our understanding of the modulating impacts of the gut microbiome on normative animal behavior.

## Introduction

There is a recent increase in studies detailing the composition of the animal gut microbiota and their influence on host behavior mediated via metabolic and biochemical linkages (Mohanta et al., 2020). Most of these studies are mainly correlative and speculative regarding these functions, with a few empirically determined ones. In essence, the gut microbiota is linked to modulating a variety of responses ranging from the animal immune system, growth, health, and behavior in many animals (De Palma et al., 2015; Davidson et al., 2018; Nagpal and Cryan, 2021; Shoji et al., 2023). This modulating effect is proposed to proceed via the vagus nerve and is mediated by microbe-derived metabolites (such as histamine, catecholamine regulators, and serotonin). These act as chemical transmitters between the gut and the brain, stimulating endocrine receptors and ultimately impacting mood and behavior (Sandhu et al., 2017; Soares et al., 2019; Mohanta et al., 2020; Williams et al., 2020; Nagpal and Cryan, 2021), in a complex and complicated cascade collectively referred to as the microbiota-gut-brain axis (MGB axis). In many animal taxa, studies demonstrate that gut microbiota is linked to exploratory behavior, neophobia, sociality, stress, and anxiety-related behaviors (Hoban et al., 2016; Burokas et al., 2017; Davidson et al., 2018; Nagpal and Cryan, 2021). However, these studies show that there is still a lot to be uncovered regarding the influence of the microbiota on the MGB axis.

Significant work with vertebrates detailing the influence of the microbiota on the MGB axis usually involves correlations between various non-typical behaviors, such as depression- and anxiety-like behaviors, and the presence of or absence of bacteria, which are then interpreted as suggestive of an effect of the gut microbiota (Nagpal and Cryan, 2021). For example, several correlative studies using fecal microbiota transplant (FMT) studies have found depression-like behaviors in recipient antibiotic-treated mice (Leclercq et al., 2020), recipient germ-free mice (Zheng et al., 2016), recipient naive mice getting FMT from vulnerable (meek) mice compared to resilient (strong) mice (Pearson-Leary et al., 2020), and in mice deficient in segmented filamentous bacteria (SFB), but reversed when gavaged with SFB noncolonized feces exhibited antidepressant behaviors(Medina-Rodriguez et al., 2020). Overall, it is difficult to assess the actual impacts of gut microbial manipulations on behavioral responses in animal models. This is due to the reliance on the emergence of “atypical” relative to “typical” behavioral responses in treated and controlled animal subjects as the best indicator of such microbial impacts.

Having well-characterized behavioral and physiological phenotypes observed and determined from selectively bred lines gives a unique opportunity to investigate the extent to which behaviors are influenced by associated gut microbiota. However, such studies are limited. Glover et al. (2021) uncovered no significant differences in fecal microbiota composition (with and without antibiotic treatment) nor an associated change in underlying behavior in low novelty responder (LR) and high novelty responder (HR) rats selectively bred to exhibit timid non-exploratory and bold and exploratory behaviors, respectively. Similarly, Suhr et al. (2023) did not detect significant differences in two distinct genetic Rainbow trout lines. In contrast, significant differences in caecal microbiomes were determined between selectively bred resilient (high litter size) and non-resilient (low litter size) rabbit lines (Casto-Rebollo et al., 2023) and dogs from well-established aggressive, phobic, or standard lines (Mondo et al., 2020). However, some evidence indicates that this can depend on whether a particular animal exists in social groups or is solitary (Pfau et al., 2023). We argue in this work that investigating the gut-brain axis and its impacts more definitively on animal behavior broadly requires the use of selectively bred lines with already established behavioral phenotypes (empirically underscored by differing neurophysiological mechanisms) rather than the use of “atypical” behaviors following treatment conditions.

Second to the mouse as a model system for studying the vertebrate MGB dynamic is the zebrafish, *Danio rerio* (Fetcho et al., 2008). Most work on the MGB in zebrafish has focused on loosely defined behavioral responses (if at all), the correlations between these in the presence of added bacteria (so-called probiotic bacteria) or the absence of bacteria (usually via antibiotic treatment) between control and treatment groups. These have ranged from decreased shoaling behavioral displays (Borrelli et al., 2016), reduced appetite (Falcinelli et al., 2016) to decreased “anxiety-like” (Davis et al., 2016) and reduced bottom-dwelling behavior (Valcarce et al., 2020), and no observed differences in “anxiety-like” behaviors (Schneider et al., 2016) between adult zebra fish fed the probiotic *Lactobacillus* relative to controls. Ironically, clearance or reduction of gut microbial diversity via antibiotic treatment also impacts the same zebrafish behavior displays and adds to the MGB phenomena. For instance, exposure to low concentrations of the antibiotic β- dike-tone increased individual exploratory behavior and group shoaling behavior but induced anxiety-like behaviors in individuals and decreased shoaling behavior at higher concentrations (Wang et al., 2016). Finally, emerging studies are utilizing gnotobiotic zebrafish larvae to elucidate neurobehavioral development. However, the observed inconsistencies (host strain used, days post-infection, husbandry condition, etc.) in the results using GF larvae emerging from these systems pose a significant challenge (Nagpal and Cryan, 2021). Thus, overall, the presence or absence of bacteria in zebrafish (because of treatment with probiotics or antibiotics) and the subsequent deviations after that from a “typical” behavioral state poses limitations on justification for associated gut microbial effects in the MGB paradigm. In contrast, given their utility as vertebrate models in models in the MGB paradigm, studies characterizing the underlying gut microbiota of selectively bred lines of zebrafish with already established behavioral phenotypes may offer new insights into this phenomenon.

To this end, we believe that animals selectively bred to display distinct and correlated suites of behavioral and physiological responses across contexts and time (i.e. personality types, stress coping styles) represent an ideal context in which to examine the MGB dynamics and whether these different phenotypes are underscored by different gut microbiota. Two common animal personality types across taxa are the bold and shy personality types. Individuals with a bold personality type are characterized by having higher exploratory and aggressive activity, and lower neophobic and glucocorticoid stress responses compared to individuals with shy personality types (Sih et al., 2004; Øverli et al., 2007; Koolhaas et al., 2010). In zebrafish, identification of bold and shy personality types have ranged from behavioral screenings of wild and lab populations to artificial selection (Baker et al., 2017). Wong et al., (2012) described the production of two selectively bred lines of zebrafish from wild caught animals, where the lines show differences in behavior consistent with the shy (HSB) or bold (LSB) personality types across 6 different behavioral assays. The differences in exploratory and stress-related behaviors between the lines are consistent across both contexts and time (Wong et al., 2012; Baker et al., 2018; Johnson et al., 2020). These two phenotypes are underscored by distinct morphology (Kern et al., 2016), basal neurotranscriptomic states (Wong and Godwin, 2015; Wong et al., 2015c), neuromolecular responses to drugs (Wong et al., 2013; Goodman and Wong, 2020), cortisol release rates in response to an acute stressor (Wong et al., 2019), and contextual fear learning and memory performances (Baker and Wong, 2019a). The behavioral differences between zebrafish personality types have also been observed in other strains of zebrafish (Bellot et al., 2022; Rajput et al., 2022; dos Santos et al., 2023).

To examine whether the cataloged differences between the selectively bred bold (HSB) and shy (LSB) lines of zebrafish are further underscored by different gut microbiota, we sequenced and characterized the associated gut microbiota of both males and females from each line. We predict that the gut microbiota are essential modulators of host behaviors within the MGB context and that the different phenotypes (shy and bold) would be underscored by distinct gut microbiome profiles (α-diversity and β-diversity). If, on the other hand, zebrafish have a stable and core microbiome assembled through dispersal and host-selective processes (Roeselers et al., 2011), one anticipates no differences in either α-diversity or β-diversity between phenotypes, suggestive of limited gut microbial control or regulation of these personality types within the MGB paradigm in this species.

## Materials and Methods

### Animal subjects

We used zebrafish from the HSB and LSB selectively bred lines (Wong et al., 2012) that show behavioral, neuroendocrine, and neuromolecular responses consistent with the shy and bold personality types, respectively. As such for simplicity, we will refer to the lines as shy and bold zebrafish. Fish were housed in mixed-sex tanks (40L) on a recirculating system with solid and biological filtration. Fish experienced a 14:10 L/D cycle with a water temperature of 26⸰C. All fish were fed twice daily with Tetramin Tropical Flakes (Tetra, Blacksburg, VA, USA). Bold (2 females and 8 males, n=10) and shy fish (3 females and 7 males, n=10) were randomly captured from their home tanks, quickly decapitated, and bodies stored at −20C until tissue processing. All fish were between 2-3 years old and had undergone 12-14 generations of selective breeding. All procedures were approved under UNO IACUC 17-070-09-FC.

### DNA extraction and microbiome sequencing

The entire digestive tract of individuals was dissected out following surface sterilization and under sterile conditions. Briefly, fish were washed for 1 minute in a 1:10 diluted detergent solution to kill any bacteria on the surface and rinsed twice for 1 minute each in nanopore water. Following manufacturer protocol, DNA was extracted from the dissected gut using the QIAGEN DNeasy PowerSoil Pro Kits (QIAGEN, Valencia, CA, USA) from the dissected gut. Extracted DNA was sequenced at the University of Nebraska Medical Center Genomics Core Facility, following high-throughput paired-end Illumina MiSeq library preparation. Briefly, a PCR reaction was performed on samples generating a single amplicon spanning the V4 (515-F) and V5 (907-R) variable region (Keskitalo et al., 2017). Library validation and DNA quantification were carried out using the Agilent BioAnalyzer 2100 DNA 1000 chip (Agilent), and Qubit 3.0 (Qubit™, Thermofisher), respectively. Pooled libraries were loaded into the Illumina MiSeq at 10 pM and spiked with 25% PhiX (a bacteriophage) for MiSeq run quality as an internal control (Mukherjee et al., 2015) to generate 300 bp paired ends with the 600-cycle kit (version 3). The raw reads were deposited into the Sequence Read Archive database (BioProject Number: PRJNA1070623).

### Data processing and statistical analyses

The R package DADA2 (version 1.26.0) was used to process fastq primer-trimmed MiSeq paired-end reads obtained from the sequencing center, phix sequences were removed, and forward and reverse reads were truncated to 290 and 280 base pairs, respectively, with median scores above 30. A naive Bayes taxonomy classifier was employed to classify each amplicon sequence variant (ASVs) against the SILVA 138.1 reference database and used to construct the taxonomy table (Wasimuddin,2020). The ASV count and taxonomy files were combined to generate a standard ASV table, filtered for sequences identified as chloroplasts, mitochondria, unassigned at the kingdom level, and eukaryotes. Further analyses were carried out in QIIME v.1.8 (Caporaso et al., 2010; Kuczynski et al., 2012; Bolyen et al., 2018).

The ASV table was summarized at the family level, and all subsequent analyses were carried out using this table. Before analyses, two samples with low reads from each group were removed, and the remaining samples were rarefied to 110 reads per sample and replicated 100 times to capture diversity (Weiss et al., 2017; McKnight et al., 2019; Cameron et al., 2021). To investigate bacterial diversity, we calculated the chao1 (Huang and Zhang, 2013), Simpson’s index (Simpson, 1949), and Shannon’s evenness (Shannon C.E, 1957) indices in QIIME. Significant differences among categorical groupings were determined using the non-parametric Wilcoxon tests in JMP Pro 15 (S.A.S., Cary, NC, USA). For compositional diversity, we generated the Bray-Curtis dissimilarity distance matrix (Bray and Curtis, 1957) using the rarefied table. This was then used to calculate non-metric multidimensional scales (NMDS)(Rabinowitz, 1975) to visualize differences in microbiome composition between behavioral phenotypes. Subsequently, differences among behavioral phenotypes were examined using permutational multivariate analysis of variance (PERMANOVA) (Anderson, 2017) with the Bray-Curtis distance matrix as input. Significant differences in the abundance of ASVs between behavioral phenotypes were examined using the group_significance command in QIIME at P < 0.05. To assess different potential metabolic /function gene profiles between the two phenotypes, we used FAPROTAX for annotation prediction(Louca et al., 2016). Significant differences in the abundance of annotated functional predicted profiles between behavioral phenotypes were examined using the group_significance command in QIIME at P < 0.05.

## Results

Quality processing (denoising, filtering, removal of phix, merging of reads, and removal of chimeras) retained 20.1% of reads (709,722 out of 3,531,286). ASV determination yielded a 1064 ASV across 20 samples. Subsequent curation of the ASV table resulted in a final filtered table of 706 ASV across 18 samples (two dropped due to low number of reads) (Num samples: 18, Num observations: 706, Total count: 65,201, with a distribution of Min: 113.000, Max: 14,907.000, Median: 1,601.500, Mean: 3,622.278, Std. dev.: 4,571.296).

An examination of unique bacterial taxa present in the gut microbiome (α-diversity) did not uncover any significant differences across the four indices examined between the bold and shy behavioral phenotypes (observed ASVs, T-test statistic: 34, P-value: 0.83), (Chao1, T-test statistic: 24.5, P-value: 0.45), (Shannon’s evenness, T-test statistic: 40, P-value: 0.41), and (Simpson’s index, , T-test statistic: 31, P-value: 0.35)(Fig. 1). We uncovered no significant sex-specific differences across behavioral phenotypes (observed ASVs, T-test statistic: 18.5, P-value: 0.95), (Chao1, T-test statistic: 22, P-value: 0.78), (Shannon’s evenness, T-test statistic: 18, P-value: 0.90), and (Simpson’s index, T-test statistic: 18, P-value: 0.90).

**Figure 1.**
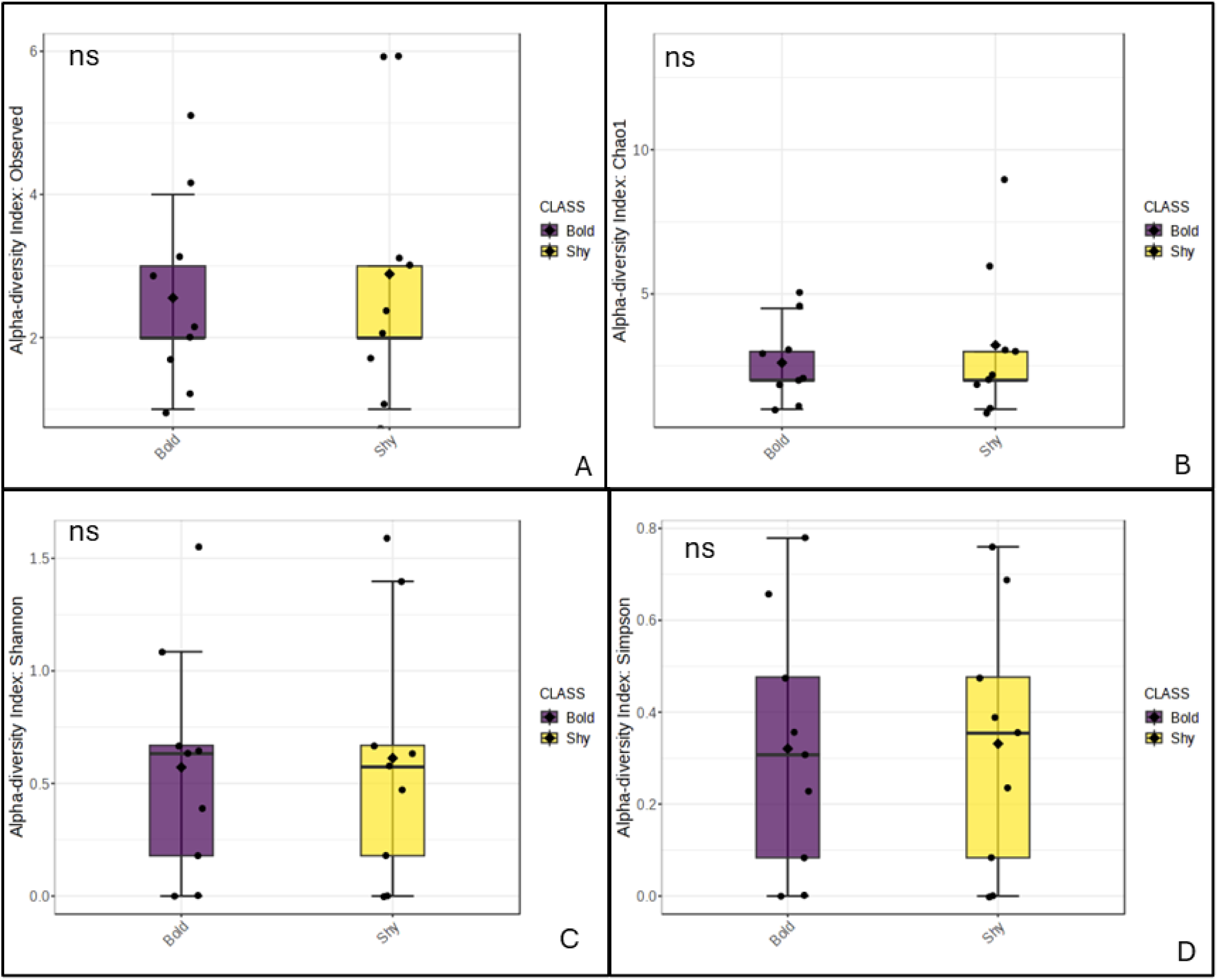
Non-significant alpha diversity estimates **A**) observed_ASVs, **B**) Chao1, **C**) Shannon’s evenness, and **D**) Simpson’s Index, between the gut microbiomes of bold (proactive) and shy (reactive) zebrafish behavioral phenotypes.

Similarly, examination of the community composition of the gut microbiomes (β-diversity) between the two behavioral phenotypes did not yield any significant differences (PERMANOVA; F-value =0.75; R^2^=0.0448; P-value=0.56) (Fig 2A). A dendrogram examining microbiome community compositions between the two did not reveal any cluster associated with behavioral phenotypes (Fig. 2B). However, no sex specific differences in microbial community composition were uncovered between the behavioral phenotypes (PERMANOVA; F-value = 0.90; R^2^ = 0.0600; P-value= 0.403).

**Figure 2.**
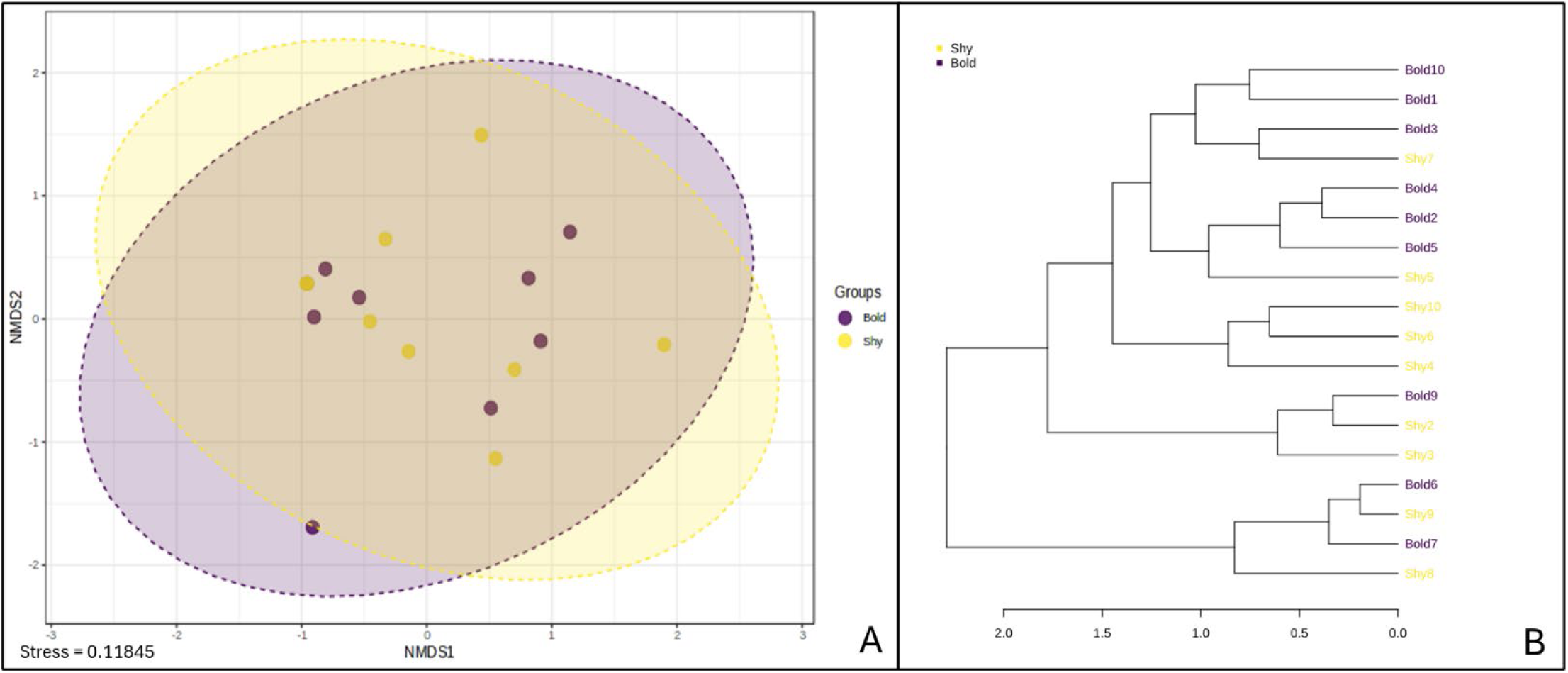
Examination of gut microbiome community composition of bold and shy zebrafish behavioral phenotypes displayed as **A**) an NMDS plot and **B**) as a dendrogram showing the absence of behavior-based clustering. (PERMANOVA; F-value =0.75; R^2^=0.0448; P-value=0.56).

Overall, core microbiome analyses revealed the presence and abundance of ∼ 16 bacterial families shared between the two behavioral phenotypes (Fig. 3A). These are bacterial taxa in both behavioral phenotypes. These 16 bacterial families are distributed across six phyla, namely, Actinomycetota (families *Myobacteriacceae* and *Streptomycetoceae*), Bacillota or Firmicutes (family *Streptococcaceae*), Bacteroidota (family *Chitiniphagaceae*), Fusobacteriota (family *Fusobacteriaceae*), Planctomycota (family *Pirellulaceae* and *Gemmataceae*), and Pseudomonadota (families *Alcaligenaceae, Aeromonadaceae, Enterobacteriaceae, Pseudomonadaceae, Rhodocbacteriaceae, Rhizobiales, Rhizobiaceae*, and *Sphingomonadaceae*). An analysis of bacterial families differentially abundant between shy and bold behavioral phenotypes (group_significance) yielded eight bacterial families at the P-value = 0.05 (Fig. 3B) (Table S1). These bacteria taxa may either be absent or present in significantly lower relative abundance in one group or the other and differ fundamentally from members of the “core” microbiota. Bacterial taxa differentially abundant in the shy zebrafish are the Pseudomonadota (Proteobacteria) (*families Beijerinckiaceae and Rhizobiales_Incertae_sedis*) and Planctomycetota (families *Pirellulaceae* and *Rubinisphaeraceae*). In contrast, the bacterial taxa Pseudomonadota (families *Burkholderiaceae, Micropepsaceae,* and *Rhodonobacteraceae*) and Actinomycetota (family *Propionibacteraceae*) are differentially abundant in the bold zebrafish. (Fig. 3B). Functional annotation based on the partial 16SrRNA gene did not yield any significant difference between the two behavioral phenotypes, which may underlie the cataloged behavioral differences (Figure S1 and Table S2).

**Figure 3.**
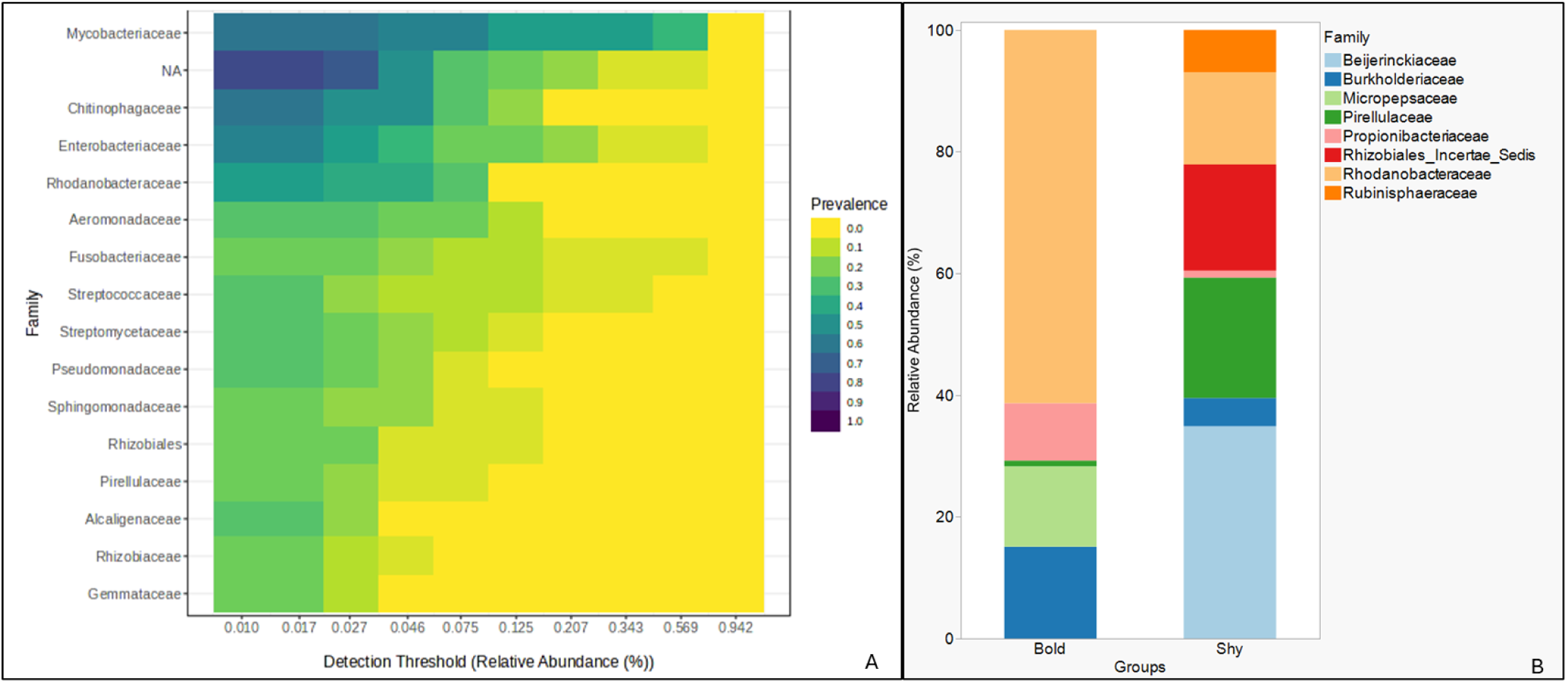
**A)** The 16 bacterial families and their relative abundances comprising the core gut microbiome of the bold and shy zebrafish behavioral phenotypes, and B) the eight differentially abundant bacterial families that vary in abundance between the bold and shy zebrafish behavioral phenotypes.

## Discussions

We characterized the gut microbiota of individuals from two distinct selectively bred lines of zebrafish that differ consistently in their exploratory behaviors and physiological responses (bold and shy personality types). Different animal behavioral phenotypes maintained and selectively bred over multiple generations may be underscored by dissimilar gut microbial community compositions. Operating within the MGB framework, we anticipated differences in gut microbiome profiles between the two distinct behavioral phenotypes. This would be underscored by different α-diversity and β-diversity measures between both phenotypes, thus highlighting microbe-mediated effects on host behavior. Alternatively, different animal behavioral phenotypes maintained and selectively bred over multiple generations may not differ significantly in community composition, suggestive perhaps of the existence of a stable “core” gut microbiome and, thus, a limited role for the gut microbiota in modulating host behavior within the MGB paradigm. The absence of significant differences in the number of unique ASVs (α-diversity) and community composition following the characterization of the gut microbiota in adult shy and bold zebrafish was unexpected in this study. Previous studies using less defined and characterized zebrafish behavioral responses have uncovered significant differences in gut microbiome composition between treatment and control adult zebrafish. In these studies, animals selectively fed with a probiotic or an antibiotic exhibited altered gut microbiome profiles, and these were associated with a behavior change. For example, significant increases in Firmicutes were reported in adult zebrafish fed the probiotic L*actobacillus rhamnosus*, resulting in decreased shoaling behavior (Borrelli et al., 2016) and reduced appetite (Falcinelli et al., 2016) relative to controls. However, the observed increase in Firmicutes in the mentioned studies is unsurprising as *Lactobacillus* fed to the treatment zebrafish is in the phylum Bacillota (formerly Firmicutes). Furthermore, although no such increases in Firmicutes were observed in zebrafish fed the probiotic, *Lactobacillus plantarum*, there were, however, limited increases in the abundances of several bacterial taxa between treatment (with reduced anxiety-like behaviors) and control individuals (Davis et al., 2016). In contrast, we used animals with inherently different behavioral phenotypes in this study. Thus, we uncovered no comparable enrichment of Firmicutes in this study, which is in contrast with studies that have found Firmicutes to be one of the dominant members of the adult zebrafish gut microbiome (Kanther and Rawls, 2010; Roeselers et al., 2011; Stephens et al., 2016; Murdoch and Rawls, 2019). It is unclear if this may be related to the two behavioral phenotypes used in this study. As far as we know, this is the only study we are aware of to characterize the *in situ* gut microbial community composition of any bold and shy zebrafish phenotypes (Bellot et al., 2022; Rajput et al., 2022) in general or of the particular genetic background from the shy and bold personality type lines (Wong et al., 2012).

In this study, the lack of dissimilarity between the two zebrafish behavioral phenotypes is supported by other zebrafish intestinal microbiota characterization studies but without a behavioral phenotype context. For example, no differences in microbiome composition were determined between wild-caught and laboratory-maintained zebrafish colonies (from multiple locations)(Roeselers et al., 2011), nor between co-housed wild-type and immune-deficient myd88 knockouts zebrafish (Burns et al., 2017). The emerging takeaway from both studies is that the zebrafish gut microbiome might be underscored by dispersal-related microbial traits, which results in a higher within-host microbial diversity but reduced overall between-host diversity (Burns et al., 2017). The reported reduced β-diversity from across these studies, ostensibly, might be indicative of a host-dependent screening or selective process that selects for a “core” associated gut microbiome despite limited variation across several laboratory-maintained zebrafish populations in multiple labs (Roeselers et al., 2011). However, the presence of a stable core gut microbiota in this species, irrespective of different behavioral phenotypes, does not suggest the absence of a modulating effect of the gut microbiota on host behavior within an MGB context. This is because the underlying premise of this study that different behavioral phenotypes would be underscored by different gut microbiota is well supported by previous studies in mice (McGaughey et al., 2019; Agranyoni et al., 2021) and by the various ways gut microbiota are postulated to modulate host behaviors.

It is important to note that it is uncertain if this study’s shy or bold behavioral phenotypes represent a dysbiotic state. While many studies compare regular to dysbiotic individuals in examining the correlations between behavior (disease state) and gut microbiota, in this study, we are not constrained to nor limited in this way, as both phenotypes can be considered “normal” and healthy. Given the well-characterized behavioral, morphological, physiological, and neurobiological differences between the shy and bold zebrafish phenotypes used in this study (Wong et al., 2012, 2019; Wong and Godwin, 2015; Kern et al., 2016; Baker et al., 2018; Baker and Wong, 2019b, 2021; Johnson et al., 2020), and despite the lack of any significant differences in potential metabolic functional profiles between the phenotypes (Fig.S1 and Table S2), it is possible that the microbiome could still be modulating the host behavior even without an underlying difference in community composition. This is true for social animals (primate and non-primates) that vary significantly in terms of within-group individual behaviors (Archie and Tung, 2015; Pasquaretta et al., 2018) but tend to have a more homogenized within-group gut microbiota (Lax et al., 2014; Moeller et al., 2016; Raulo et al., 2021).

Gut microbes modulate animal behavior within the MGB context by producing metabolites (or their precursors) that function as chemical communication signals between the gut and the nervous and endocrine systems (Schretter, 2020). Short-chain fatty acids (SCFAs) produced by a plethora of fermentative gut-associated bacteria in animals (Silva et al., 2020), as well as other microbe-produced neurotransmitters, are known to influence behaviors (Homer et al., 2023). Dopamine, acetylcholine, serotonin, and gamma-aminobutyric acids (GABA) are some examples of neurotransmitters demonstrated to be synthesized both by the neurons and by some gut bacteria (Wong et al., 2015a; Silva et al., 2020; Homer et al., 2023). Members of the phylum Actinomycetota, particularly *Bifidobacterium,* produce GABA, which influences behaviors. Similarly, Propionibacteriaceae (phylum Actinomycetota), which was abundant in Bold zebrafish in this study, produces propionate, an essential SCFA (Turgay et al., 2022) that may be involved in modulating this behavioral phenotype in bold relative to shy zebrafish. However, several members of other bacterial phyla determined to be differently abundant in this study (Pseudomonadota, Planctomycetota, and Actinomycetota) in both shy and bold zebrafish are known SCFA-producing taxa (Deleu et al., 2021; Frolova et al., 2022), making it challenging to assign differences between these two zebrafish lines to bacterial taxonomy and abundance. The possibility remains, however, that the differentially abundant taxa (even in the absence of significant dissimilarity between the two phenotypes) may be mediating processes related to observed differences in physiological markers, such as cortisol (Wong et al., 2019), memory (Baker and Wong, 2019), and neurotranscriptomic expressions (Wong et al., 2015) between these two lines.

In conclusion, the results of our study suggest that behaviorally distinct and cataloged zebrafish phenotypes are not underscored by statistically significant differences in gut microbiome diversity and composition. This starkly contrasts with studies utilizing disruption or supplementation approaches to modulating the gut microbiome and examining the impact of these treatments on animal behaviors. In these studies, the “response” behaviors are not always as well characterized as the intrinsic behavioral phenotypes in this study. The implications of the results in this study for gut microbe-mediated behavioral responses within the MGB paradigm are unclear. However, as a first step, utilizing well-characterized and cataloged behaviors in gut microbiome disruption or supplementation studies in the MGB context might be a more rigorous experimental approach to yield empirical data supporting the mediator effects of gut microbiota on animal behavior.

## Acknowledgments

This study was funded by the University of Nebraska, Omaha, start-up grant to Dr. Paul Ayayee. This work was also completed using the University of Nebraska DNA Sequencing Core, which receives partial support from the National Institute for General Medical Science (NIGMS) INBRE-P20GM103427-19 grant and the Fred & Pamela Buffett Cancer Center Support Grant-P30 CA036727. Financial support for husbandry and maintenance of the zebrafish used in this study was provided by the University of Nebraska at Omaha’s Animal Care and Use Program and National Institutes of Health R15MH113074 and IDeA under Grant # 5P20GM103427 to RYW. We acknowledge the assistance of various undergraduate lab assistants in the Ayayee Microbial Ecology Group for assistance with the dissection and extraction of DNA and the animal husbandry staff and Wong lab members for maintaining the zebrafish facility.

## Ethical Approval

The Institutional Animal Care and Use Committee at the University of Nebraska-Omaha approved all procedures involving animals (protocol # UNO IACUC 17-070-09-FC).

## Competing interests

The authors declare no competing or financial interests.

## Data Availability Statement

The authors confirm that the data supporting the findings of this study are available within the article and its supplementary materials.

## Declarations

### Ethics approval and consent to participate

Not applicable.

### Consent for publication

Not applicable.

### Competing interests

The authors declare no competing interests.

## Author Contributions

PAA and RYW conceived and designed the study. RYW initiated and housed the zebrafish. PAA prepared samples for processing, and PAA analyzed data. PAA and RYW wrote the submission.

## Notes

### Competing Interest Statement

The authors have declared no competing interest.

