## supplementary data for "Zebrafish (*Danio rerio*) behavioral phenotypes not underscored by different gut microbiota"

**Supplementary Figure 1.** Relative abundance of top 30 functional ecological microbial cluster genes that varied between the bold and shy behavioral phenotypes following FAPROTAX analysis.

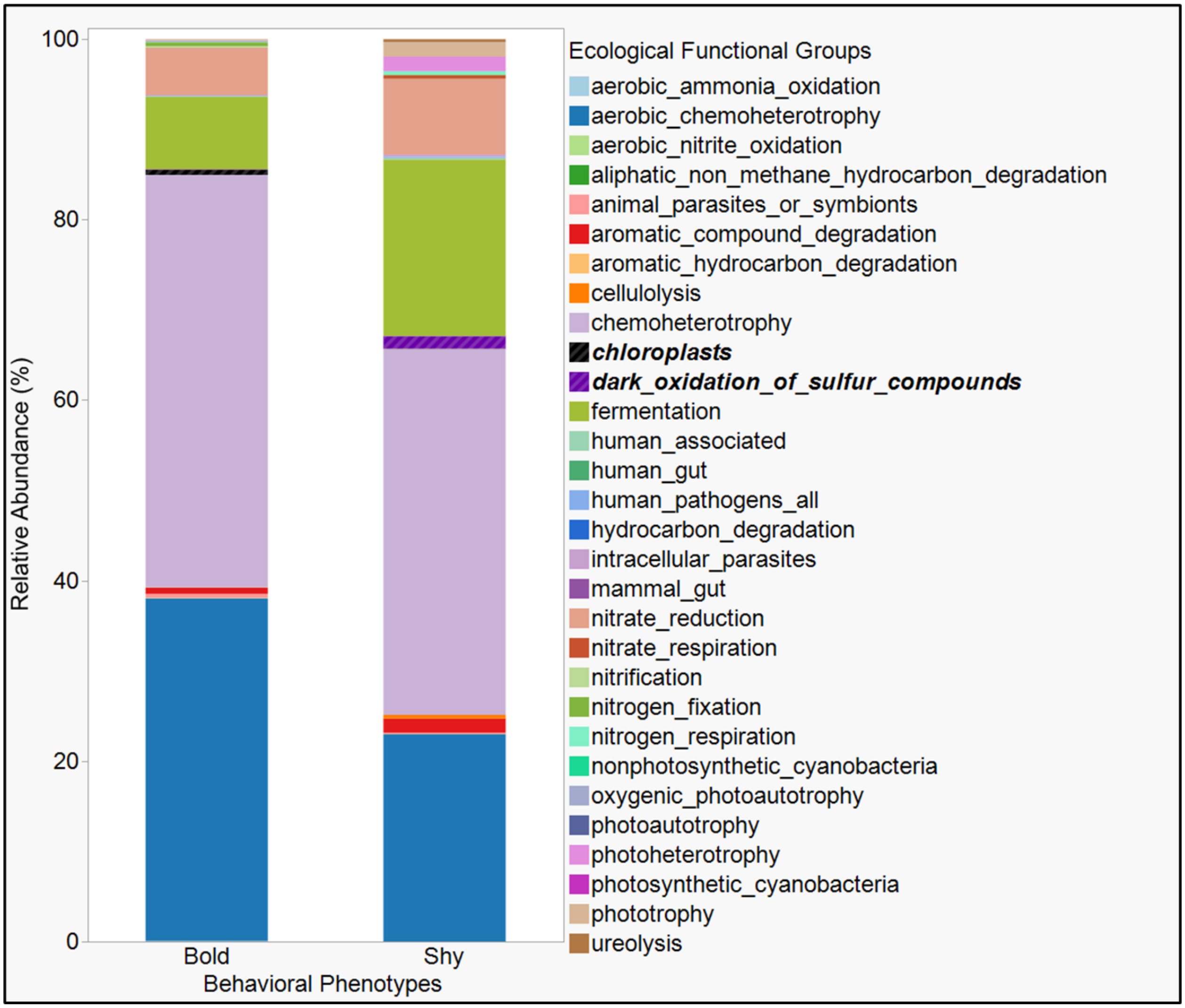

**Supplementary Table 1**. Relative abundances of the eight differentially abundant bacterial phyla and families between the bold and shy zebrafish behavioral phenotypes

| Phylum | Family | Shy | Bold |
|---|---|---|---|
| Pseudomonadota | Beijerinckiaceae | 34.88372 | 0 |
| Pseudomonadota | Rhodanobacteraceae | 15.11628 | 61.32075 |
| Planctomycetota | Rubinisphaeraceae | 6.976744 | 0 |
| Pseudomonadota | Micropepsaceae | 0 | 13.20755 |
| Pseudomonadota | Rhizobiales_Incertae_Sedis | 17.44186 | 0 |
| Pseudomonadota | Burkholderiaceae | 4.651163 | 15.09434 |
| Planctomycetota | Pirellulaceae | 19.76744 | 0.943396 |
| Actinomycetota | Propionibacteriaceae | 1.162791 | 9.433962 |

**Supplementary Table 2**. Output of different functional ecological microbial cluster genes that varied between the bold and shy behavioral phenotypes. The OTU-style functional gene profile first table was converted to a biom file and the script group_significance.py in qiime was executed on the resulting biom table. The functional ecological group chloroplast and dark_oxidation_of_sulfur_compounds were marginally significant (P = 0.05 and 0.06, respectively).

| Functional Group | Test-Statistic | P | Shy_mean | Bold_mean |
|---|---|---|---|---|
| ***chloroplasts*** | ***3.76*** | ***0.05*** | ***1.56*** | ***13.78*** |
| ***dark_oxidation_of_sulfur_compounds*** | ***3.45*** | ***0.06*** | ***140.11*** | ***1.56*** |
| human_pathogens_all | 3.01 | 0.08 | 13.78 | 0.89 |
| human_associated | 3.01 | 0.08 | 13.78 | 0.89 |
| chemoheterotrophy | 2.39 | 0.12 | 4044.56 | 1077.89 |
| nitrogen_fixation | 2.14 | 0.14 | 7.33 | 10.89 |
| aerobic_chemoheterotrophy | 2.12 | 0.15 | 2290.67 | 895.78 |
| photoheterotrophy | 2.12 | 0.15 | 161.00 | 1.89 |
| cellulolysis | 1.62 | 0.20 | 39.56 | 0.78 |
| ureolysis | 1.62 | 0.20 | 32.44 | 0.89 |
| phototrophy | 1.47 | 0.23 | 161.00 | 2.11 |
| intracellular_parasites | 1.34 | 0.25 | 21.89 | 1.00 |
| aerobic_nitrite_oxidation | 1.00 | 0.32 | 0.00 | 1.33 |
| aromatic_hydrocarbon_degradation | 1.00 | 0.32 | 0.44 | 0.00 |
| aliphatic_non_methane_hydrocarbon_degradation | 1.00 | 0.32 | 0.44 | 0.00 |
| hydrocarbon_degradation | 1.00 | 0.32 | 0.44 | 0.00 |

| | | | | |
|---|---|---|---|---|
| photosynthetic_cyanobacteria | 1.00 | 0.32 | 0.00 | 0.22 |
| oxygenic_photoautotrophy | 1.00 | 0.32 | 0.00 | 0.22 |
| photoautotrophy | 1.00 | 0.32 | 0.00 | 0.22 |
| aerobic_ammonia_oxidation | 0.83 | 0.36 | 1.67 | 1.33 |
| nitrate_respiration | 0.70 | 0.40 | 36.89 | 1.44 |
| nitrogen_respiration | 0.70 | 0.40 | 36.89 | 1.44 |
| animal_parasites_or_symbionts | 0.65 | 0.42 | 14.78 | 10.56 |
| fermentation | 0.56 | 0.45 | 1950.11 | 189.22 |
| nonphotosynthetic_cyanobacteria | 0.37 | 0.54 | 0.44 | 1.00 |
| nitrate_reduction | 0.16 | 0.69 | 835.89 | 122.89 |
| nitrification | 0.03 | 0.87 | 1.67 | 2.67 |
| aromatic_compound_degradation | 0.01 | 0.93 | 155.22 | 15.89 |
| human_gut | 0.01 | 0.94 | 1.67 | 0.89 |
| mammal_gut | 0.01 | 0.94 | 1.67 | 0.89 |
| methanotrophy | nan | nan | 0.00 | 0.00 |
| acetoclastic_methanogenesis | nan | nan | 0.00 | 0.00 |
| methanogenesis_by_disproportionation_of_methyl_groups | nan | nan | 0.00 | 0.00 |
| methanogenesis_using_formate | nan | nan | 0.00 | 0.00 |
| methanogenesis_by_CO2_reduction_with_H2 | nan | nan | 0.00 | 0.00 |
| methanogenesis_by_reduction_of_methyl_compounds_with_H2 | nan | nan | 0.00 | 0.00 |

| | | | | |
|---|---|---|---|---|
| hydrogenotrophic_methanogenesis | nan | nan | 0.00 | 0.00 |
| methanogenesis | nan | nan | 0.00 | 0.00 |
| methanol_oxidation | nan | nan | 0.00 | 0.00 |
| methylotrophy | nan | nan | 0.00 | 0.00 |
| sulfate_respiration | nan | nan | 0.00 | 0.00 |
| sulfur_respiration | nan | nan | 0.00 | 0.00 |
| dark_sulfite_oxidation | nan | nan | 0.00 | 0.00 |
| sulfite_respiration | nan | nan | 0.00 | 0.00 |
| thiosulfate_respiration | nan | nan | 0.00 | 0.00 |
| respiration_of_sulfur_compounds | nan | nan | 0.00 | 0.00 |
| arsenate_detoxification | nan | nan | 0.00 | 0.00 |
| arsenate_respiration | nan | nan | 0.00 | 0.00 |
| dissimilatory_arsenate_reduction | nan | nan | 0.00 | 0.00 |
| arsenite_oxidation_detoxification | nan | nan | 0.00 | 0.00 |
| arsenite_oxidation_energy_yielding | nan | nan | 0.00 | 0.00 |
| dissimilatory_arsenite_oxidation | nan | nan | 0.00 | 0.00 |
| anammox | nan | nan | 0.00 | 0.00 |
| nitrate_denitrification | nan | nan | 0.00 | 0.00 |
| nitrite_denitrification | nan | nan | 0.00 | 0.00 |
| nitrous_oxide_denitrification | nan | nan | 0.00 | 0.00 |

| | | | | |
|---|---|---|---|---|
| denitrification | nan | nan | 0.00 | 0.00 |
| chitinolysis | nan | nan | 0.00 | 0.00 |
| knallgas_bacteria | nan | nan | 0.00 | 0.00 |
| dark_hydrogen_oxidation | nan | nan | 0.00 | 0.00 |
| nitrate_ammonification | nan | nan | 0.00 | 0.00 |
| nitrite_ammonification | nan | nan | 0.00 | 0.00 |
| nitrite_respiration | nan | nan | 0.00 | 0.00 |
| xylanolysis | nan | nan | 0.00 | 0.00 |
| dark_sulfide_oxidation | nan | nan | 0.00 | 0.00 |
| dark_sulfur_oxidation | nan | nan | 0.00 | 0.00 |
| dark_thiosulfate_oxidation | nan | nan | 0.00 | 0.00 |
| manganese_oxidation | nan | nan | 0.00 | 0.00 |
| manganese_respiration | nan | nan | 0.00 | 0.00 |
| ligninolysis | nan | nan | 0.00 | 0.00 |
| invertebrate_parasites | nan | nan | 0.00 | 0.00 |
| human_pathogens_septicemia | nan | nan | 0.00 | 0.00 |
| human_pathogens_pneumonia | nan | nan | 0.00 | 0.00 |
| human_pathogens_nosocomia | nan | nan | 0.00 | 0.00 |
| human_pathogens_meningitis | nan | nan | 0.00 | 0.00 |
| human_pathogens_gastroenteritis | nan | nan | 0.00 | 0.00 |

| | | | | |
|---|---|---|---|---|
| human_pathogens_diarrhea | nan | nan | 0.00 | 0.00 |
| fish_parasites | nan | nan | 0.00 | 0.00 |
| plant_pathogen | nan | nan | 0.00 | 0.00 |
| oil_bioremediation | nan | nan | 0.00 | 0.00 |
| dark_iron_oxidation | nan | nan | 0.00 | 0.00 |
| iron_respiration | nan | nan | 0.00 | 0.00 |
| fumarate_respiration | nan | nan | 0.00 | 0.00 |
| chlorate_reducers | nan | nan | 0.00 | 0.00 |
| predatory_or_exoparasitic | nan | nan | 0.00 | 0.00 |
| anoxygenic_photoautotrophy_H2_oxidizing | nan | nan | 0.00 | 0.00 |
| anoxygenic_photoautotrophy_S_oxidizing | nan | nan | 0.00 | 0.00 |
| anoxygenic_photoautotrophy_Fe_oxidizing | nan | nan | 0.00 | 0.00 |
| anoxygenic_photoautotrophy | nan | nan | 0.00 | 0.00 |
| aerobic_anoxygenic_phototrophy | nan | nan | 0.00 | 0.00 |
| plastic_degradation | nan | nan | 0.00 | 0.00 |
| reductive_acetogenesis | nan | nan | 0.00 | 0.00 |
